# ST2 Signaling Regulates Innate Immune Responses in Kidney Injury

**DOI:** 10.64898/2026.01.24.701541

**Authors:** Vikram Sabapathy, Gabrielle Costlow, Saanvi Acharya, Clint Upchurch, Oliver Pelletier, Bushra Mekhri, Timothy Bullock, Norbert Leitinger, Sanja Arandjelovic, Rahul Sharma

## Abstract

**Introduction:** Innate immune cells are critical in inflammation, repair, and fibrosis post-kidney injury. Nuclear-cytokine interleukin (IL)-33, which is released upon tissue damage, signals through IL-1-receptor-like-1 (IL1RL1 or ST2), expressed on many immune cells, including macrophages. However, macrophage regulation by IL-33/ST2 is incompletely understood. We hypothesized that ST2 plays a vital role in activating and/or mobilizing myeloid cells and macrophages to sites of injury.

**Methods:** We performed acute and chronic ischemia-reperfusion injury (IRI) in mice with myeloid cell-specific deletion of ST2 (ST2^fl/fl^.LysM^Cre^) to examine the role of myeloid cells ST2 expression in renal injury. The structure and function of the kidney were probed using flow cytometry, histology, immunohistochemistry, quantitative gene expression, and biochemical analysis. The *invitro* efferocytosis assay, RNA Seq, and Seahorse assay were carried out using bone-marrow-derived macrophages

**Results:** Interestingly, ST2 deletion resulted in attenuated renal pathology in the acute renal IRI model, whereas in chronic IRI, the loss of ST2 exacerbated kidney injury, suggesting a role of ST2 in the resolution of chronic injury. RNA sequencing (RNASeq) analysis of bone-marrow-derived ST2 sufficient and deficient macrophages showed that loss of ST2 downregulated genes involved in oxidative phosphorylation and clearance of dead cells (efferocytosis). Indeed, the ST2-deficient macrophages had reduced phagocytosis activity. Further, Seahorse analysis revealed that ST2-deficient macrophages had compromised mitochondrial metabolism.

**Conclusions:** We conclude that the IL-33/ST2 axis is essential for regulating macrophage function and contributes to regulating tissue homeostasis following renal injury.

## 1. Introduction

Acute kidney injury (AKI) is a life-threatening disease affecting 10-15% of hospitalized patients and 50% of patients in intensive care units (ICU) worldwide and is currently without effective early diagnosis or targeted therapy^1^. AKI also predisposes to chronic kidney disease (CKD) that leads to end-stage renal disease (ESRD). Kidney-related ailments result in around 2 million patients succumbing to the disease worldwide due to the shortage of donor kidney availability^2,3^.

Ischemic injury of the kidneys results in tubular cell death leading to the clinical syndrome of acute tubular necrosis. Postmortem examination of human specimens has shown that ischemic tubular injury leads to an accumulation of inflammatory mononuclear cells in the vasa recta of the outer medulla^4^. Similarly, animal models of ischemia-reperfusion also demonstrate that renal ischemic injury (IRI) is characterized by the rapid influx of polymorphonuclear leukocytes, lymphocytes, and macrophages into the renal interstitium^5,6^. Innate immune cells play a critical role in the development and resolution of AKI^7^. Innate immune cells such as macrophages and neutrophils are activated in response to AKI, producing cytokines and chemokines that can also cause tissue damage^8^. This innate immune response is initiated within minutes of renal injury^9^. The kidney resident macrophages occupy specific niches in rodent and human kidneys, and in response to injury attain a developmental state and unique distribution^10,11^.

On the other hand, the adaptive immune response, which is mounted at later points (days) after the injury occurs, is antigen-dependent and targeted. Innate and adaptive immune responses are not mutually exclusive mechanisms of host defense but rather synergetic^12^. Defects in either system results in immune dysregulation. Cells involved in innate immune response include macrophages, neutrophils, dendritic cells, mast cells, basophils, eosinophils, natural killer (NK) cells, and innate lymphoid cells (ILCs). The rapid response of these cells is mediated through production of alarmins, cytokines and chemokines^13^, which further potentiate cell recruitment and regulate the inflammatory milieu.

Phagocytic actions of macrophages also promote the clearance of damaged cells, cell debris, and pathogens from the site of injury or inflammation^14^. Studies have also suggested that boosting efferocytosis in macrophages could assist in resolving inflammation^15^. In addition, macrophages present antigens to T cells^16^, thus acting as important messengers between innate and adaptive immunity. Macrophages obtain distinct phenotypes depending on the physiological conditions in response to stimulation^17^. Generally, macrophages are classified into classically activated (also referred to as “M1”) or alternatively activated (“M2”) macrophages. M1 macrophages are characterized by the release of pro-inflammatory mediators through engagement with T helper 1 (Th1) cells. Contrastingly, M2 macrophages promote immunoregulatory signals through their cooperation with T helper 2 (Th2) cells^18^. Molecular mechanisms involved in the plasticity and transitions between these macrophage states have been reported to have conflicting results with M2 macrophages being contributors to protection from or resolution of AKI as well as contributing to fibrosis during CKD^19^. Thus, the molecular mechanisms of macrophage polarization during ischemic injury are not completely understood.

The alarmin cytokine interleukin 33 (IL-33), a member of the IL-1 cytokine family, is constitutively expressed in epithelial cells, fibroblasts, and endothelial cells and is released as a consequence of tissue injury. Upon release, IL-33 binds to the ST2 receptor expressed on most myeloid cells, such as macrophages, mast cells, eosinophils, basophils, and natural killer cells (reviewed in)^20^. ST2 exists in two splice variants^21^: membrane-bound ST2, which signals via downstream activation of MyD88/NF-κB and p38 MAPK pathway, and soluble ST2 (sST2), which acts as a decoy receptor and sequesters free IL-33. Short-term activation of naïve macrophages with IL-33 has been shown to promote their transition to the M1 phenotype; however, prolonged activation elicits the transition of M1 macrophages to the M2 phenotype^22,23^.

In this study, we asked whether ST2 signaling in myeloid cells determines the outcome of acute and chronic ischemia-reperfusion injury in mice and whether ST signaling in macrophages is required for their homeostatic functionality.

## 2. Materials and Methods

### 2.1. Animal Models

All the animal experiments in the study were approved by the University of Virginia Animal Care and Use Committee (ACUC) in accordance with the NIH Guide for the Care and Use of Laboratory Animals. The *IL1Rl1*^fl/fl^ (*IL1RL1*^tm1a^) mice were obtained from the UCDAVIS KOMP Repository (KOMP Project ID CSD35155), USA. *LysM^Cre^* (*LysM^Cre^*(*Lyz2*-Cre^24^, Jackson Laboratory, Strain #004781) mice were purchased from Jackson Laboratory (Bar Harbor, USA). *IL1Rl1^fl/fl^ LysM^Cre^* was generated by crossing *IL1Rl1*^fl/fl^ and *LysM^Cre^* mice. Primers used for genotyping are listed in **Supplementary Table 3**.

### 2.2. Cell Culture

The bone marrow-derived macrophages were isolated as previously described^25^. The macrophages were cultured in DMEM supplemented with 10% fetal bovine serum (FBS) and 10% L929 conditioned media.

### 2.3. Renal Function

The mice were anesthetized with ketamine-xylazine (i.p.), and blood was collected using the heparinized capillary tube retro-orbitally. The blood plasma separated from whole blood was used to measure plasma creatinine (PCr) and blood urea nitrogen (BUN). The plasma creatinine (PCr) was measured enzymatically per manufacturer instructions with minor modifications in which the sample volume was doubled (Diazyme Laboratories). BUN was quantitated using a kit (Arbor Assay) as per manufacturer instructions.

### 2.4 Histology and evaluation of kidney injury and fibrosis

Kidney sections stained with Hematoxylin and Eosin (H&E) were used to evaluate kidney injury and scored as previously described^26^. Kidney fibrosis was histologically assessed using Masson’s Trichrome staining and quantified using Image J.

### 2.5. Immunofluorescence

Kidney sections were fixed with 1% paraformaldehyde (PFA) for 72 hours and embedded in an optimal cutting temperature (OCT) compound (Ted Pella, Inc.). The tissue was sectioned 5μm thick, permeabilized with 0.2% Triton X-100, and blocked using 1% FBS. The samples were first incubated with the unconjugated antibody overnight at 4°C. Following primary antibody incubation, fluorescent-labeled secondary antibody was added and incubated at room temperature for 4 hours. All the samples were mounted using ProLong Diamond Antifade (Thermo Fischer Scientific) containing 4’,6-diamidino-2-phenylindole (DAPI), a nuclear counterstain. Microscopic images were acquired using the Carl Zeiss Axiovert 200 microscope system with Apotome Image Resolution and Axiovision software (Carl Zeiss Microscopy, LLC).

### 2.6. Apoptosis Analysis

Apoptosis in kidney tissue was quantified using a fluorescein *In Situ* Cell Death Detection Kit based on the TUNEL principle (ROCHE) per the manufacturer’s instructions.

### 2.7. Real-time gene expression analysis

RNA was isolated using the RNeasy kit (Qiagen, Germany), and the complementary DNA (cDNA) was prepared using the iScript cDNA synthesis kit (BioRad). Real-time gene expression analysis of samples was carried out using iQ SYBRGreen Supermix (BioRad) in CFX Real-Time PCR Detection System (BioRad). The gene expression levels were normalized using the glyceraldehyde 3-phosphate dehydrogenase (*Gapdh*) housekeeping gene. Primers used in QPCR are listed in Supplementary Table 2.

### 2.8. Flow cytometry

Flow cytometry was performed as previously described^27,28^. Antibodies used for the analysis are listed in the **Supplementary Table 1**. Cytek Northern Lights (Cytek Biosciences, CA) 3 laser spectral flow cytometer was used to acquire the samples. FlowJo software (BD Biosciences, NJ) was used for data analysis. The gating strategies used to analyze kidney-specific Neutrophils, Macrophages, and Dendritic Cells are shown in **Supplementary Figure 7**. For gating strategy for kidney Tregs and T cell cytokine expression refer to earlier study^26^, **Supplementary Figure 3** and **Supplementary Figure 5**.

### 2.9. Efferocytosis Assay

Human Jurkat T cells were stained with CypHer5E (GE Healthcare, PA15401) for 30 minutes at 37°C water bath. The labeled cells were subjected to apoptosis with 150mJ/cm^2^ ultraviolet C irradiation and incubated in a 5%CO_2_ incubator for 4 hours at 37°C. The levels of apoptosis were confirmed using Annexin V and 7AAD staining by flow cytometry. Macrophages and apoptotic cells were incubated in a 1:10 ratio for 1 hour. Following incubation, the coculture media was washed, and macrophages were dissociated using trypsin for downstream analysis using a flow cytometer. A group containing cytochalasin D was used as a control.

### 2.10. Metabolic Flux Analyzer (Seahorse) Assay

The mitochondrial stress test (MST) was performed as previously described^29^. The macrophages (1×10^4^) were seeded into the 96-well seahorse tissue culture plate (Agilent Technologies) and cultured overnight. Prior to assay, UV-irradiated Jurkat cells were seeded at a 1:10 ratio and incubated for 1 hour. Following incubation, the wells were washed and fresh media DMEM with pyruvate (Thermo-Fisher, Cat#:12800017; pH = 7.35 at 37°C) was added and equilibrated for 30 min^29^. Oxygen consumption rate (OCR) from the cell media was measured using Seahorse XF96 Flux Analyzer (Agilent Technologies). After three basal OCR measurements, compounds to modulate cellular respiratory function [1μM Oligomycin (Sigma-Aldrich); 2μM BAM15 (Cayman Chemical Company); 1μM Antimycin A and 100 nM Rotenone (Sigma-Aldrich)] were injected into the plate, and cellular respiratory rates were measured. Experiments and analysis were carried out as per the manufacturer’s instructions.

### 2.11. RNA sequencing

RNA was extracted from cells using the RNeasy Plus mini kit (Qiagen). The purity and quantity of the RNA were measured using Nanodrop 2000 (Thermo Scientific) and HS-RNA Tape Station (Agilent Technologies). The ribosomal RNA (rRNA) was then removed from the total RNA before preparing the transcriptome library. The RNA sequencing was performed using the NGS Nextseq kit – 150 cycle High Throughput kit (Illumina), with 75bp paired-end sequencing on the Illumina Nextseq 2000 sequencing system. The quality of the library preparation was checked using FASTQC to evaluate the quality of the sequencing quality and data analysis performed as before^30^. Briefly, the fastq files were trimmed to remove the Illumina adapter sequences, and low-quality sequencing reads using cutadapt with a phred quality cutoff of 20, then reassessed using FastQC. The trimmed sequencing reads were aligned to the mouse reference genome mm10 using HISAT2. FeatureCounts was used to count the number of reads mapped to genes, creating a count matrix. Differential gene expression was analyzed using DESeq2. The gene expression was visualized using a principle component analysis and heatmap. Gene ontology and Kyoto Encyclopedia of Genes and Genomes (KEGG) pathway analysis were performed using goseq^31^. The full dataset can be found in the National Center for Biotechnology Information (NCBI) Gene Expression Omnibus (Geo) database, with an Accession Number: GSE288154 (Reviewer Access Token: mhghwaimdhulhqx)

### 2.12. Statistical Analyses

All the statistical analyses for the experiments were performed in GraphPad Prism10. The wild-type (WT) and mutant alleles were analyzed using a one-way analysis of variance (ANOVA), followed by Tukey’s post hoc test for multiple comparisons. The results are presented as the mean ± standard error of the mean (SEM), with *P*<0.05 considered significant.

## 3. Results

### 3.1. Loss of ST2 in Myeloid Cells Attenuates Acute Renal Injury

To test if ST2 is expressed in murine renal macrophages, we performed a Python programming-based meta-analysis of single-cell RNA seq data from kidney resident CD45^+^ cells (Zenodo: 7314511) using Scanpy ^32,33^. Among the various immune cell subsets (**Figure 1A**), we were able to identify robust ST2 expression in kidney tissue-resident macrophages (**Figure 1B**).

**Figure 1:**
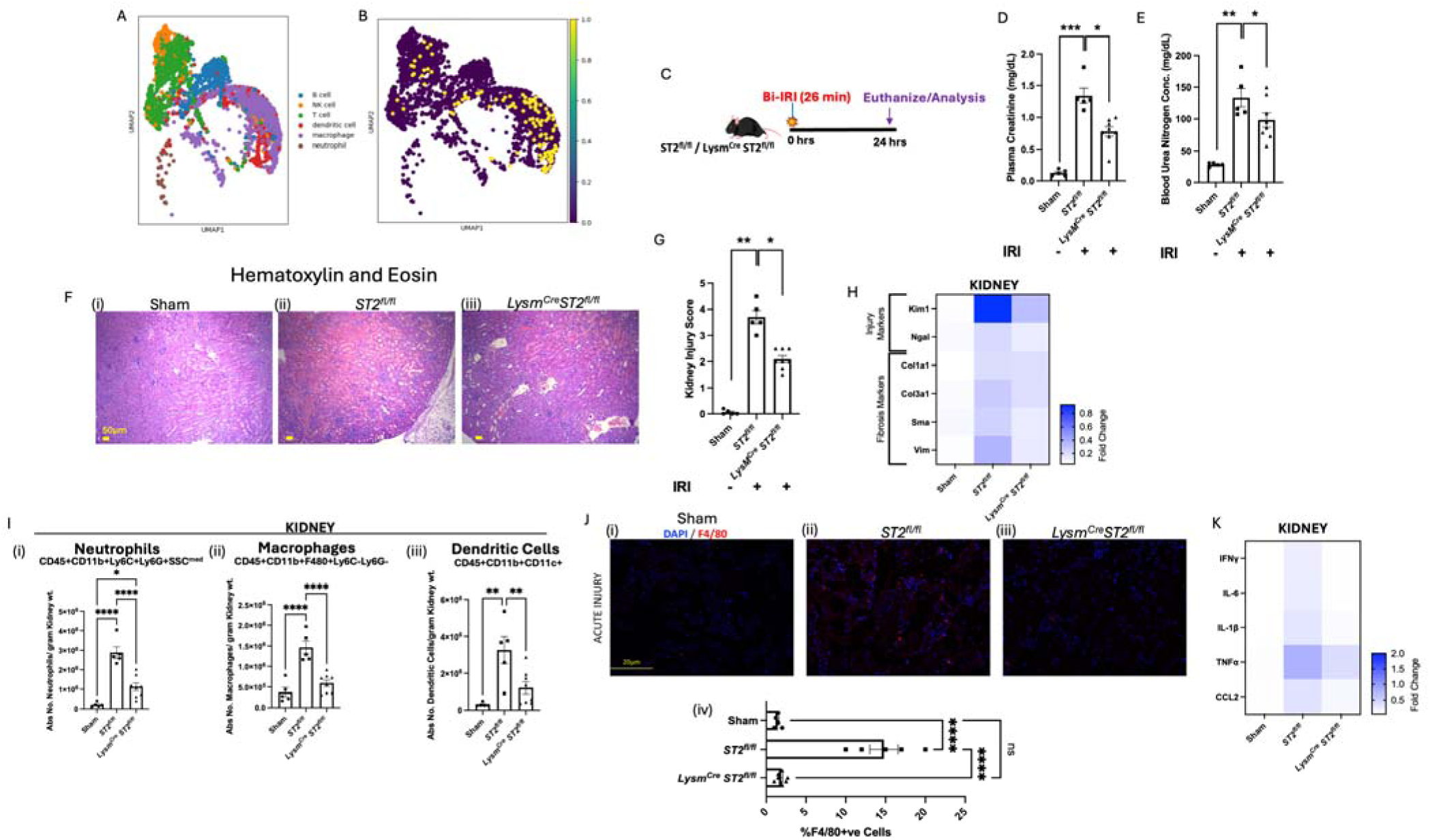
Absence of ST2 signaling in myeloid cells during acute phase attenuates AKI by lowering innate immune response and inflammation. (A) Meta-analysis of single-cell RNAseq ata: UMAP clustering of flow-sorted CD45^+^ immune cells from wild-type kidney. (B) Feature plot of *St2* expression: Visualization of *St2* expression across immune cell populations, indicating that its expression is primarily localized to macrophages. (C) Schematic representation of AKI model: Ischemia-reperfusion injury (IRI) induced in *ST2^fl/fl^*and *ST2^fl/fl^ LysM^Cre^* mice by bilateral renal pedicle clamping for 26 minutes. Kidneys were harvested 24 hours post-IRI for downstream analysis. Plasma creatinine (D) and BUN (E) results demonstrate that the loss of ST2 in myeloid cells resulted in the reduction of acute injury compared to their wild-type counterparts. (F) The histological examination revealed significantly less tubular damage in *ST2^fl/fl^ LysM^Cre^* (iii) mice compared to *ST2^fl/fl^* mice(ii); Scale bar = 50μm. (G) Kidney Injury Score: Quantitative scoring of tubular injury from histological sections, showing significant reduction in *ST2^fl/fl^ LysM^Cre^* mice compared to *ST2^fl/fl^* (H) Heatmap representing real-time quantitative gene expression of renal injury (Kim1 and Ngal) and fibrosis markers (Col1a1, Col3a1, Sma, and Vim) suggests ST2 deficiency in myeloid cells reduces the expression of injury and fibrosis-associated genes. (I) Flowcytometry analysis quantifying the presence of (i) Neutrophils, (ii) Macrophages, and (iii) Dendritic cells in the kidney following AKI reveals significant attenuation of myeloid cells infiltrating the *ST2^fl/fl^ LysM^Cre^* mice kidneys; (J) Quantification of immunofluorescence staining for F4/80+ cells confirms significant reduction in macrophage infiltration in *ST2^fl/fl^ LysM^Cre^* mice; Scale bar = 20μm. (K) Quantitative real-time analysis of proinflammatory cytokines and Chemokines (IFNγ, TNFα, IL-6, IL-1β, and CCL2) in kidneys. Symbols represent individual mice; n≥5, mean±SEM is shown; *p<0.05; **p<0.01; ***p<0.001; ****p<0.0001.

Next, to understand the role of ST2 in myeloid cell-mediated immunity, we bred *ST2^fl/fl^* mice with *LysM^Cre^* mouse line to generate mice lacking ST2 expression in myeloid cells (*ST2^fl/fl^ LysM^Cre^*mice). We then subjected *ST2^fl/fl^ LysM^Cre^* and *ST2^fl/fl^* (control) mice to a model of ischemia-reperfusion-induced acute kidney injury (IRI-AKI) (**Figure 1C**). Ischemia was induced by the clamping of the renal artery for 26 min before reperfusion was allowed. Mice were euthanized after 24 hours of reperfusion, and plasma and kidney tissue were collected for analysis. Loss of ST2 in myeloid cells resulted in significant attenuation of acute renal injury, as evidenced by reduced levels of plasma creatinine (**Figure 1D**) and blood urea nitrogen (BUN, **Figure 1E**) in *ST2^fl/fl^ LysM^Cre^* mice, compared to *ST2^fl/fl^*controls. Analysis of IRI-AKI histological samples showed significantly less tubular damage in the *ST2^fl/fl^ LysM^Cre^* mice compared to *ST2^fl/fl^* mice **(Figure 1F, 1G)**. Whole kidney quantitative gene expression analysis indicated that the kidney injury markers Kim1 and Ngal were significantly downregulated in *ST2^fl/fl^ LysM^Cre^* when compared to *ST2^fl/fl^* mice. Similarly, expression of the renal fibrosis markers *Col1a1, Col3a1, Sma*, and *Vim* was reduced in *ST2^fl/fl^ LysM^Cre^* mice (**Figure 1H**, **Supplementary Figure 4**).

To understand how loss of ST2 in myeloid cells influences inflammatory infiltration, we performed flow cytometry analysis of the IRI-AKI renal tissue. The accumulation of myeloid cells including neutrophils, macrophages, and dendritic cells in the kidneys of IRI-AKI *ST2^fl/fl^*controls was dramatically reduced in *ST2^fl/fl^ LysM^Cre^* mice (**Figure 1I**). Reduction of F4/80^+^ macrophages in IRI-AKI *ST2^fl/fl^ LysM^Cre^* mice, was also observed by immunofluorescence staining of F4/80^+^ cells as compared to *ST2^fl/fl^* mice (**Figure 1J**). Gene expression analysis revealed reduced transcripts for IFNγ, TNFα, IL-6, IL-1β, and Ccl2 in the IRI-AKI kidneys of *ST2^fl/fl^ LysM^Cre^* mice, compared to *ST2^fl/fl^* controls, suggestive of reduced renal inflammation in the absence of myeloid ST2 expression (**Figure 1K**).

### 3.2. Loss of ST2 in Myeloid Cells Exacerbates Renal Injury Under Chronic Conditions

We next sought to determine if the loss of ST2/IL-33 signaling in myeloid cells would alleviate long-term chronic renal pathology. *ST2^fl/fl^ LysM^Cre^* mice and *ST2^fl/fl^* controls were subjected to unilateral ischemia-reperfusion (Uni-IRI), followed by the nephrectomy of the contralateral kidney two weeks later, a day prior to euthanasia to measure the function of the injured kidney (**Figure 2A**). Surprisingly, plasma creatine and BUN values indicated that renal function was significantly deteriorated in *ST2^fl/fl^ LysM^Cre^*mice compared to *ST2^fl/fl^* mice (**Figure 2B-C**). Histological analysis also indicated higher tubular injury, glomerular sclerosis, and immune cell infiltration in mice lacking ST2 expression in myeloid cells (**Figure 2D-E**). Quantitative gene expression analysis of the renal tissue indicated that mRNA expression of kidney injury markers *Kim1* and *Ngal* was significantly increased in *ST2^fl/fl^ LysM^Cre^* when compared to *ST2^fl/fl^* mice, as were the kidney fibrosis markers *Col1a1, Col3a1, Sma*, and *Vim* (**Figure 2F**, **Supplementary Figure 5**). Masson’s trichrome staining also indicated that *ST2^fl/fl^ LysM^Cre^* mice exhibited increased fibrosis compared to *ST2^fl/fl^* mice (**Figure 2H-I**). Flow cytometry data showed no significant difference in neutrophils or dendritic cells in the kidneys of *ST2^fl/fl^ LysM^Cre^* mice compared to *ST2^fl/fl^* controls; however, macrophage numbers were significantly reduced (**Figure 2G**). Despite reduced macrophage accumulation, kidney gene expression analysis revealed increased expression of several inflammatory cytokines (IFNγ, TNFα, IL-6, IL-1β, and CCL2) in the *ST2^fl/fl^ LysM^Cre^* mice as compared to *ST2^fl/fl^* mice (**Figure 2J**). These data suggest that although the loss of ST2 in myeloid cells impairs macrophage accumulation, it contributes to elevated inflammation and renal damage in chronic renal injury.

**Figure 2:**
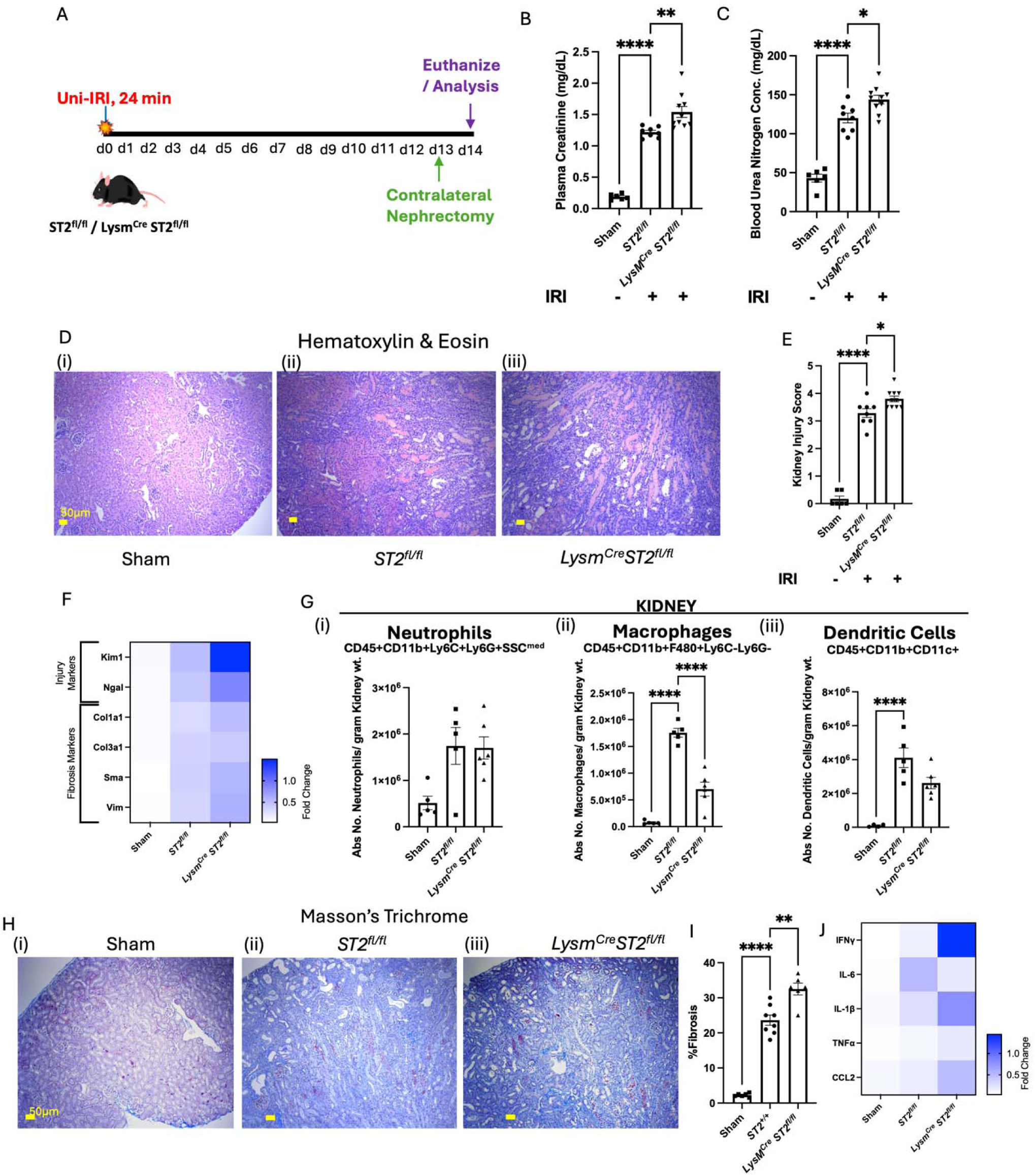
Persistent loss of ST2 signaling in myeloid cells leads to exacerbation of chronic kidney disease. (A) Schematic representation of the experimental timeline for the chronic injury model, indicating unilateral ischemia-reperfusion injury (Uni-IRI) for 24 minutes, followed by contralateral nephrectomy on day 13, a day before final euthanasia of mice on day 14. Renal functional parameters were measured on plasma samples collected on day 14 of the chronic kidney disease (CKD) model. Plasma Creatinine (B) and Blood Urea Nitrogen assay (C) levels were significantly elevated in mice lacking ST2 signaling in myeloid cells, *ST2^fl/fl^ LysM^Cre^* compared to controls (sham and *ST2^fl/fl^*). (D) Histological assessment of kidney damage using Hematoxylin and Eosin (H&E) staining; Scale bar = 50μm. The quantitative representation of tubular and glomerular damage, interstitial fibrosis, and inflammation using Kidney Injury Score (E) shows *ST2^fl/fl^ LysM^Cre^* mice had more prominent kidney injury. (F) Heatmap representing real-time quantitative gene expression of renal Injury (Kim1 and Ngal) and fibrosis markers (Col1a1, Col3a1, Sma, and Vim). (G) Flow cytometry analysis quantifying the presence of (i) Neutrophils, (ii) Macrophages, and (iii) Dendritic cells in the kidney during CKD. (H) Evaluation of renal fibrosis by Masson’s Trichrome Staining (MTS) reveals increased collagen deposition (blue staining) in injured mice; Scale bar = 50μm. (I) MTS data quantification demonstrates significantly higher fibrotic scores in *ST2^fl/fl^ LysM^Cre^* mice compared to *ST2^fl/fl^*. (J) Quantitative real-time analysis of pro-inflammatory genes (IFNγ, TNFα, IL-6, IL-1β, and CCL2) in kidney tissues at day 14. *ST2^fl/fl^ LysM^Cre^* mice displayed heightened expression of these markers, indicative of amplified inflammatory response. Symbols represent individual mice; n≥5, mean±SEM is shown; *p<0.05; **p<0.01; ***p<0.001; ****p<0.0001.

### 3.3. Macrophage – T Cell Cross Talk

In the acute IRI-AKI model, loss of ST2 in macrophages also led to reduced systemic immune activation after acute IRI-AKI, as observed by significantly reduced TNFα and IL-4 production and a trend towards lower production of IFNγ and IL-10 in CD4^+^ T-cells (**Figure 3A**) suggesting a role of ST2 expression in antigen-presenting cells affecting T-cells. Since the T-cell activation takes a few days to fully mature, we also studied the effects of loss of ST2 in myeloid cells in the chronic injury model. While there were no significant differences observed in the levels of TNFα and IFNγ production, the ability of CD4^+^ T-cells to produce IL-4 and the immunoregulatory cytokine IL-10 was markedly attenuated in the mice with ST2-deficient myeloid cells (**Figure 3C**). The data suggests that ST2 expression on myeloid-cell regulated T-cell response may play a role in IRI-AKI. Since the production of IL-10, a cytokine also produced by the regulatory T-cells (Treg), we hypothesized that macrophage cross-talk with Treg may be critical to mitigate inflammation and initiate repair processes. Therefore, we tested if Treg numbers were affected in mice with myeloid ST2 deletion during renal injury. In both acute and chronic IRI, Treg numbers were elevated in the kidneys of injured *ST2^fl/fl^* mice (**Figure 3B-D**) as compared to the sham mice. However, we noted that in chronic renal injury, Treg levels were significantly lower in *ST2^fl/fl^ LysM^Cre^* mice than in *ST2^fl/fl^* controls (**Figure 3D**), suggesting that reduced Treg numbers due to loss of ST2 expression in myeloid cells could contribute to greater chronic renal injury in these mice.

**Figure 3:**
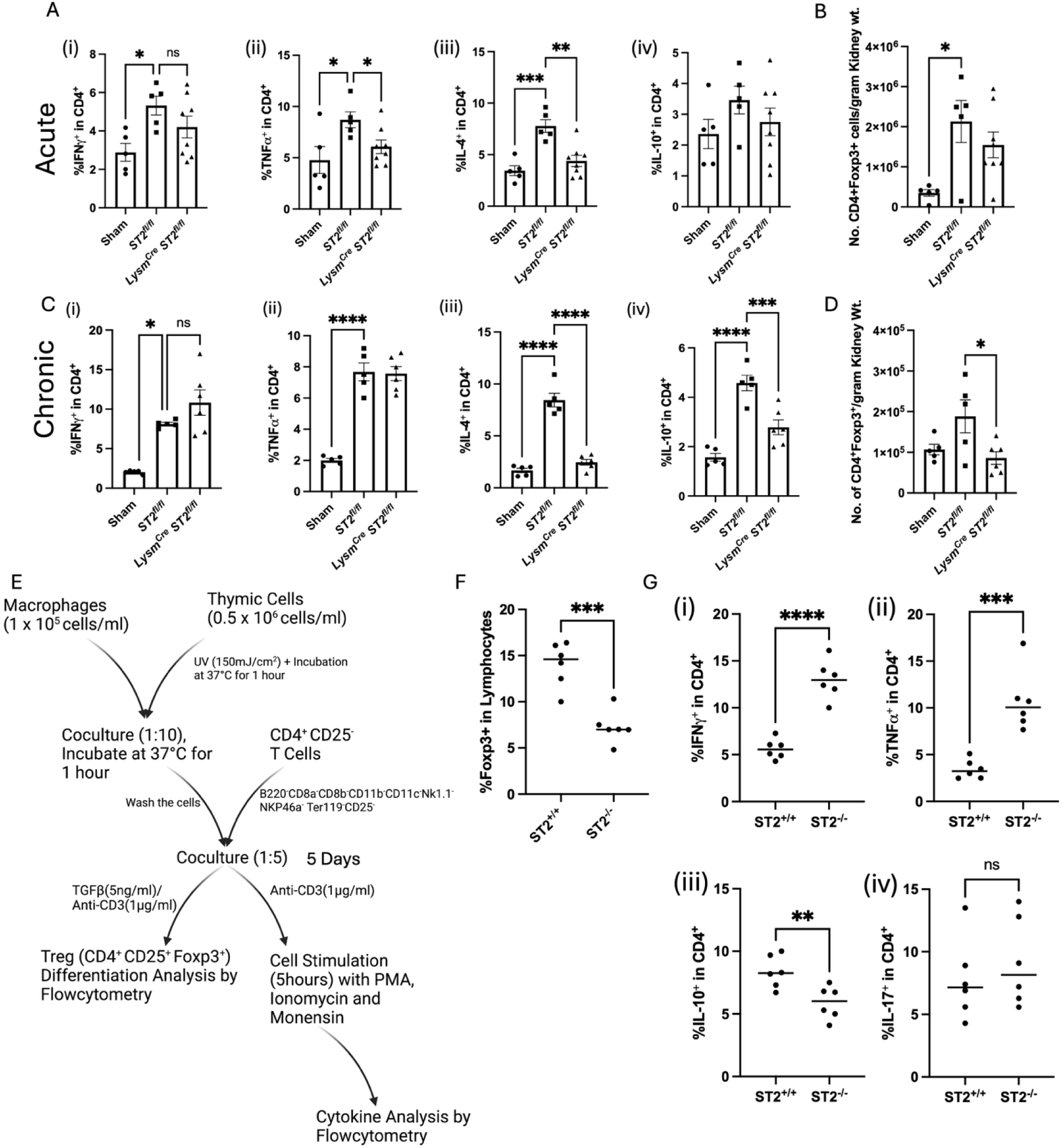
Macrophage-T Cell cross-talk is critical for Immune cell homeostasis. (A) Flow cytometry analysis of cytokines (Ifnγ, Tnfα, Il-4, and Il-10) in CD4+ Tcells during (A) acute and (C) chronic kidney injury; Flowcytomtery analysis of kidney-specific regulatory T cells (Tregs) in (B) Acute and (D) chronic kidney injury models. The results suggest significant attenuation of Tregs in ST2^fl/fl^ LysM^Cre^ mice following chronic injury. (E) Schematic representation of the in vitro coculture system: Bone marrow-derived macrophages (BMDM) and thymic cells were isolated; UV-treated apoptotic thymic cells were cocultured with BMDM for BMDM activation; Activated BMDM was cocultured with splenic CD4^+^CD25^-^ T cell in the presence of TGFβ and ant-CD3 for Treg differentiation and only anti-CD3 for cytokine analysis. (F) Flow cytometry analysis of T cell differentiation shows an increase in Tregs generation when cocultured in the presence of ST2 sufficient macrophages; (G) Proportion of CD4^+^ lymphocytes expressing IFNγ, TNFα, IL-10, and IL-17 quantified via flow cytometry following macrophage-T cell coculture. IFNγ^+^CD4^+^ T-cells/TNFα^+^CD4^+^ T-cells proinflammatory Th1 skewed lymphocytes were elevated during T cell co-culture with ST2 deficient BMDM. Symbols represent individual mice (A-D; n≥5); In vitro experiments, **Figure E-G** (two replicates of n=3); mean±SEM is shown; *p<0.05; **p<0.01; ***p<0.001; ****p<0.0001.

We next used a macrophage-T cell co-culture system to evaluate how macrophage ST2 expression affects Treg differentiation and cytokine expression (**Figure 3E**). The bone marrow-derived macrophages (BMDM) were isolated from *ST2^fl/fl^ LysM^Cre^* and *ST2^fl/fl^* mice. The isolated macrophages were activated by UV-treated wild-type thymic cells. Responder T cells were isolated from the spleens of wild-type mice. The T cells were purified using magnetic-activated cell sorting (MACS) to ensure nonregulatory (CD25-negative), highly pure CD4^+^ T cells. Co-culture assay of BMDM with responder T cells showed significantly reduced Foxp3^+^ Treg differentiation in the co-cultures of ST2 deficient macrophages (**Figure 3F**). Flow cytometry-based cytokine analysis of the responder cells indicated that the pro-inflammatory cytokines IFNγ and TNFα were significantly elevated in T cells co-cultured with ST2 deficient BMDM (**Figure 3G(i) and (ii)**), whereas the anti-inflammatory cytokine IL-10 was reduced (**Figure 3G(iii)**) and there was no significant difference in IL-17 (**Figure 3G(iv)**). Collectively, these data suggest that reduced Tregs in myeloid ST2 deficiency and increased inflammatory cytokine production could contribute to chronic renal pathology.

### 3.4. ST2 deficiency changes the transcriptional landscape in macrophages

To discern how the loss of ST2 expression influences macrophage function, we performed bulk RNAseq analysis of BMDM from ST2^-/-^ and wildtype (ST2^+/+^) controls. Bone marrow-derived monocyte differentiated BMDM were used rather than mature macrophages from tissues to understand the early events and avoid the changes accumulated throughout the lifespan due to ST2 deficiency. Euclidean distance algorithm indicated that the whole transcriptomic profile of ST2 deficient macrophages was highly distinct from control macrophages (**Figure 4A**). Heatmap of differential gene expression suggested distinct gene expression in ST2 knockout macrophages (**Figure 4B**). A volcano plot was used to identify top-upregulated and downregulated transcripts (**Figure 4C**). The gene *Bcl2a1b* helps in balancing cell survival and programmed cell death, shaping macrophage persistence^34^. It is a highly regulated nuclear factor kB (NF-κB) target gene. Arginase (*Arg1*) directs polarization of macrophages towards the anti-inflammatory M2 phenotype^35^. (Growth Differentiation Factor 15) *Gdf15*^36^ modulates the inflammatory process and plays an important role in tissue repair. *Cd38*^37^ is involved in regulating inflammatory responses and influences macrophage activation states, Chitinase-like protein-1 ^38^ (*Chil1*) modulates the secretion of pro-inflammatory factors by macrophages, such as TNFα and Monocyte chemoattractant protein-1 (MCP1) thus influencing macrophage polarization to anti-inflammatory M2 phenotype. Ly6c2^39,40^ characterizes macrophage subsets, guiding immune surveillance. *Chil3*^35^ fosters an anti-inflammatory environment by responding to damaging stimuli. Collectively, the upregulated genes in ST2 expressing macrophages influence the immunomodulatory landscape. Downstream pathway analysis showed many functional pathways such as immune cell activation, wound healing response, metabolism-associated pathways such as oxidative phosphorylation and glycolysis, chemokine response, cytokine receptor activity, and phagosome formation (**Supplementary Figure 1**) were highly influenced by ST2 expression in BMDM (**Figure 4D-F**).

**Figure 4:**
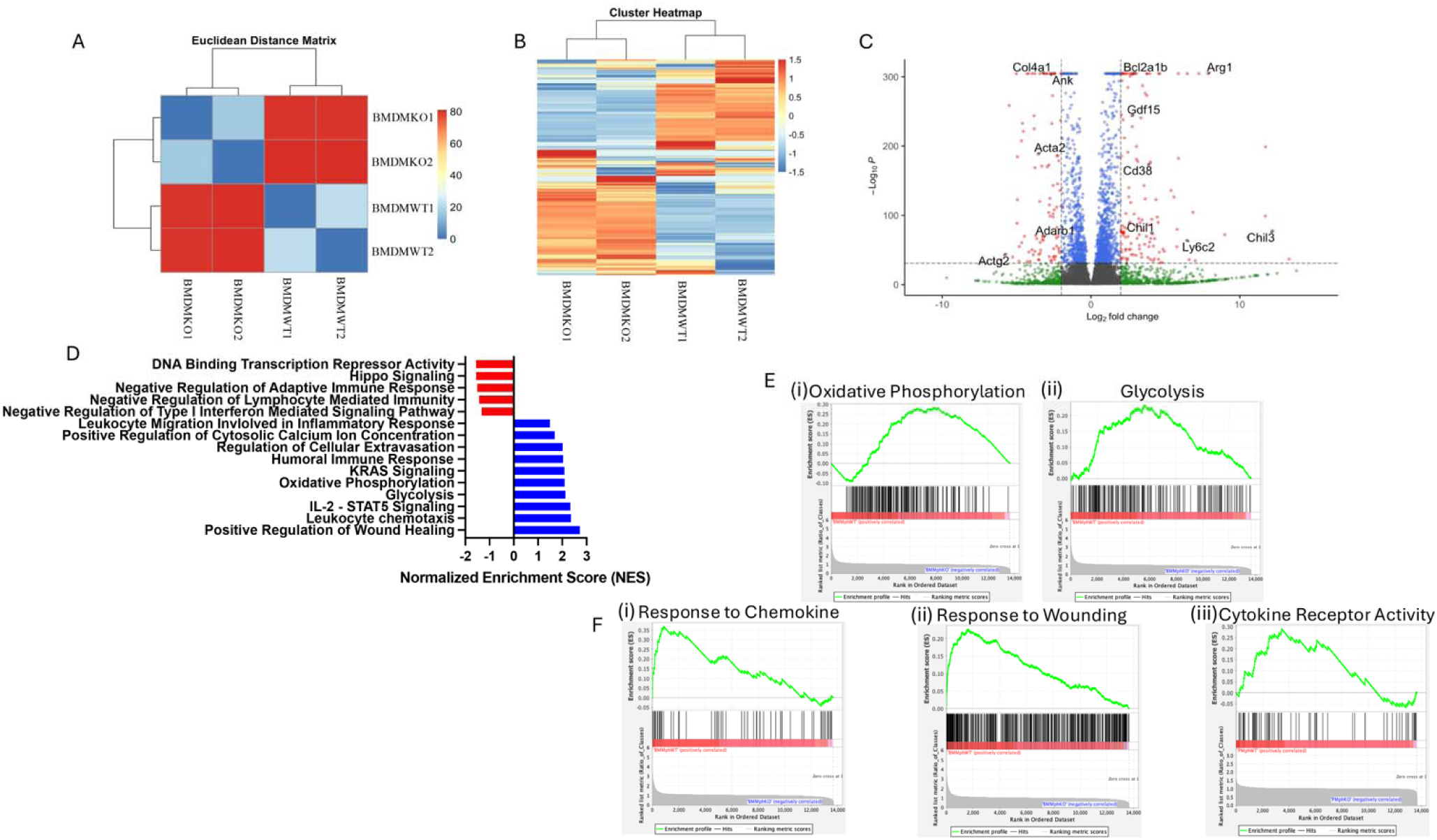
RNA sequencing analysis showing differential gene expression analysis between ST2 sufficient (BMDMWT) and deficient (BMDMKO) bone marrow derived macrophages. (A) Eucledian distant matrix indicating the distance of the cell type based on whole transcriptomics. A small distance between two cells in the matrix indicates that their gene expression profiles are very similar, suggesting they might belong to the same cell type. (B) Heatmap of differentially expressed genes (DEGs) highlights the distinct gene expression signatures between BMDMWT (n=2) and BMDMKO (n=2) macrophages; (C) Volcano plot illustrating the DEGs between BMDMWT vs BMDMKO, the x-axis represents the log2 fold change, and the y-axis represents the -log_10_P-value. ST2 sufficient macrophages exhibited upregulation of Bcl2a1b, Arg1, Gdf15, Cd38, Chil1, Chil3, and Ly6c2 immunomodulatory and reparative genes. (D) Gene ontology analysis of bone marrow macrophage showing normalized enrichment scores (NES), pathways related to Leukocyte Trafficking, Humoral Immune Response, Oxidative Phosphorylation, Glycolysis, Leukocyte chemotaxis, and Regulation of Wound Healing were markedly upregulated in BMDMWT. (E) Gene set enrichment analysis (GSEA) plot shows (i) Oxidative Phosphorylation, (ii) Glycolysis, (iii) Chemokine Response, (iv) Response to injury, and (v) Cytokine receptor activity positively correlated with BMDMWT.

### 3.5. ST2 Deficiency Impairs Normal Homeostatic functions of Macrophages

Macrophage numbers were reduced in the injured kidneys of *ST2^fl/fl^ LysM^Cre^* mice in both the acute and chronic renal injury, compared to *ST2^fl/fl^* controls (**Figures 1I(ii), J** and **2G(ii)**). Targeted analysis of genes essential for macrophage development and function showed downregulation of key markers, including the co-stimulatory molecule *CD80*, required for effective T-cell activation^41^; *Mfge8* (Milk Fat Globule-EGF Factor 8), a glycoprotein facilitating phagocytic clearance of apoptotic cells^42^; *Itgb3* (Integrin Beta 3), an adhesion molecule essential for apoptotic cell clearance and immune regulation^43^; *Abca1* (ATP-Binding Cassette Transporter A1), crucial for maintaining lipid homeostasis in macrophages^44^; and *Stab2* (Stabilin-2), a scavenger receptor involved in apoptotic cell clearance^45^ (Figure 5A). Indeed, KEGG pathway analysis also showed defective phagosome formation in ST2-deficient BMDM (**Supplementary Figure 1**). Efficient efferocytosis is critical for the resolution of inflammation and re-establishment of homeostasis after tissue injury^46^. Therefore, we tested if apoptotic cell clearance is impaired in *ST2^fl/fl^ LysM^Cre^* mice during kidney injury. TUNEL staining revealed an elevated presence of apoptotic cells in *ST2^fl/fl^ LysM^Cre^*mice compared to *ST2^fl/fl^* mice during both acute and chronic renal inflammation (**Figure 5B-C**). We next tested if the macrophage capacity to clear dying cells by efferocytosis *ex vivo* was reduced in ST2-deficient macrophages. We incubated BMDM from *ST2^fl/fl^ LysM^Cre^* and *ST2^fl/fl^* mice with apoptotic cells labeled with the pH-sensitive dye CypHer-5e (which becomes fluorescent upon acidification in the macrophage phagolysosomes), and corpse engulfment was analyzed by flow cytometry. Efferocytic activity was significantly lower in ST2-deficient BMDM than in wild-type macrophages (**Figure 5D**). As a control, we incubated macrophages with cytochalasin D to inhibit the actin cytoskeleton rearrangements, which blocked efferocytosis in both ST2 deficient and control macrophages (**Figure 5D**). These data collectively suggest that increased numbers of TUNEL-stained apoptotic cells in the kidneys of *ST2^fl/fl^ LysM^Cre^* mice could stem from decreased macrophage-mediated efferocytosis. We further evaluated if macrophage proliferation is impaired in the absence of ST2 expression. We used dehydrogenase-dependent CCK8 assay to detect live cell doubling time in *in vitro* cultured macrophages from *ST2^fl/fl^ LysM^Cre^* and *ST2^fl/fl^* mice. We noted that the doubling time of ST2 deficient macrophages was significantly higher compared to WT macrophages (**Figure 5E-F**), suggesting that the loss of the ST2 receptor in BMDM decreases the macrophage proliferation rate.

**Figure 5:**
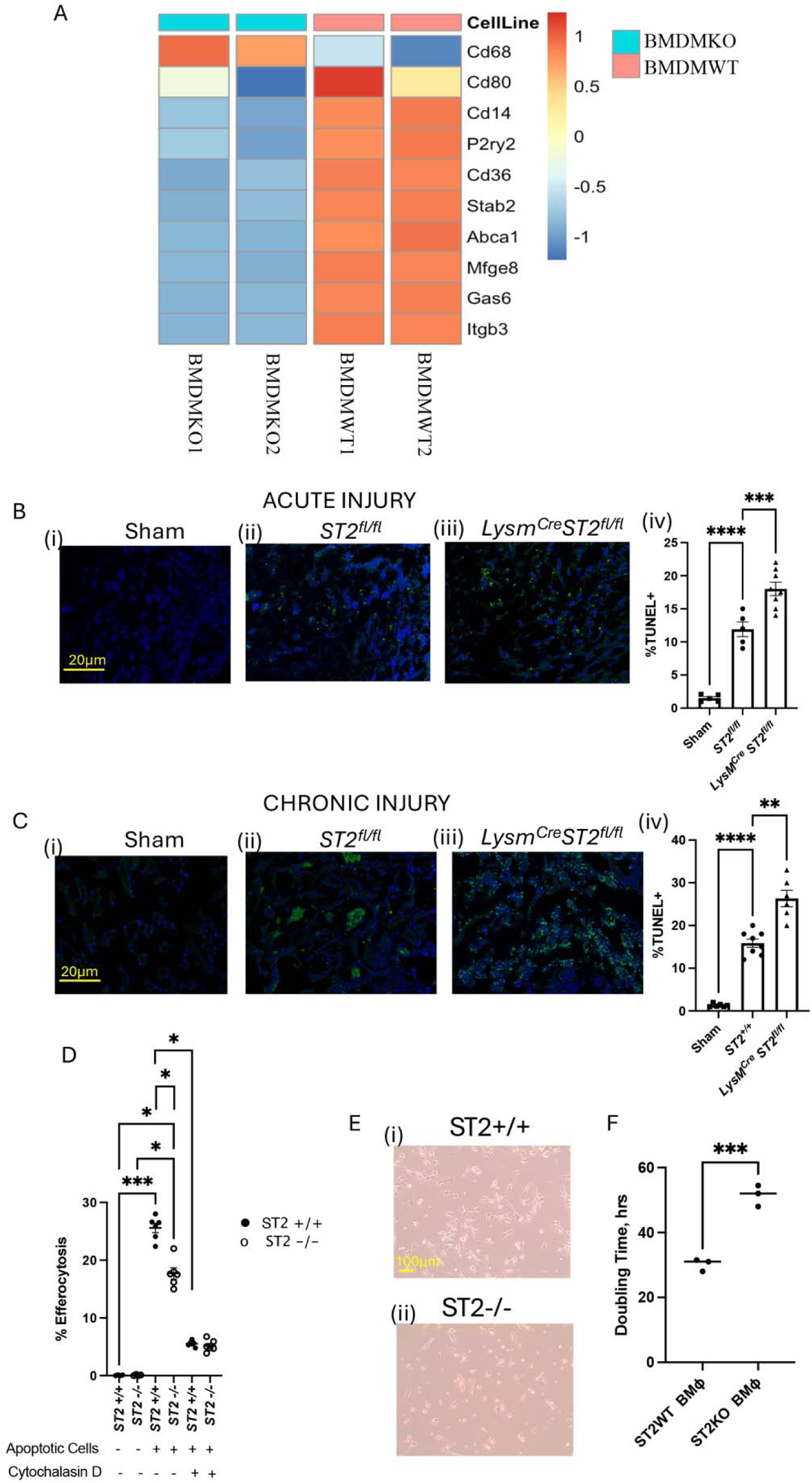
Loss of ST2 affects macrophage proliferation and efferocytosis activity. (A) Heatmap representing target gene expression profiles of macrophage-associated markers between BMDMWT (n=2) and BMDMKO (n=2). The analysis highlights significant downregulation of genes associated with macrophage development and functionality in BMDMKO. (B) Quantification of the apoptotic cells by TUNEL staining in acute (B) and chronic (C) kidney injury models; Scale bar = 20μm. Quantifying immunofluorescence staining reveals increased presence of TUNEL+ cells in *ST2^fl/fl^ LysM^Cre^* mice kidneys during the injury’s acute (Biii-iv) and chronic phases (Ciii-iv). (D) Efferocytosis activity of bone marrow-derived macrophages. The percentage of macrophages engulfing apoptotic cells was quantified, indicating impaired efferocytosis in ST2-deficient macrophages. (E) Light microscopy image of BMDMWT and BMDMKO groups, illustrating differences in cell proliferation. Representative images highlight morphological changes associated with ST2 loss; Scale bar = 100μm. (F) Quantification of Macrophage proliferation rates in WT and ST2KO groups (passage 1) demonstrates a significant reduction in cell proliferation in the absence of ST2 signaling, confirming its role in supporting macrophage growth. Symbols represent individual mice (B-C; n≥5); *In vitro* experiments, D three replicates, n=2 and F three replicates, n=1; mean±SEM is shown; *p<0.05; **p<0.01; ***p<0.001; ****p<0.0001.

### 3.6. ST2 Regulates Mitochondrial Metabolism in Macrophages

Macrophage efferocytosis and proliferation, including efferocytosis-induced proliferation, are cellular processes that are highly energy-dependent^47,48^. We next hypothesized that the macrophage metabolism could be altered in the absence of ST2 expression. To address this, we subjected ST2KO and ST2WT BMDM to Seahorse metabolic flux analyzer analysis, using the Mito Stress Test assay (Agilent). ST2KO macrophages exhibited reduced oxidative phosphorylation rates compared to control macrophages, as evidenced by a lower oxygen consumption rate (OCR) at baseline (blue vs. green bars, **Figure 6B(i)** and reduced maximal respiration (blue vs. green bars, **Figure 6B(ii)**). This highlights impaired mitochondrial energetics in ST2-deficient macrophages under homeostatic conditions. Upon exposure to apoptotic cells, OCR increased in wild-type and ST2KO macrophages. However, this increase was significantly greater in ST2-sufficient macrophages than ST2KO macrophages (red vs. magenta bars, **Figure 6B(ii)**), indicating that ST2 signaling promotes enhanced oxidative metabolism upon activation by apoptotic cells. ATP production followed a similar trend, with wild-type macrophages exhibiting higher ATP production than ST2KO macrophages at baseline (blue vs. green bars, **Figure 6B(iii)**). This difference was further amplified upon exposure to apoptotic cells (red vs. magenta bars, **Figure 6B(iii)**), underscoring the diminished metabolic flexibility of ST2-deficient macrophages. Overall, OCR changes upon activation (baseline to apoptotic conditions) were more pronounced in wild-type macrophages (**Figure 6B**, blue to red bars, compared to ST2KO macrophages (**Figure 6B**, green to magenta), suggesting impaired metabolic activation in the absence of ST2 signaling. The Interpretation of energy map data showed that macrophages with the loss of ST2 predominately remained in the quiescent state, compared to controls. During efferocytosis, the ST2-deficient BMDM used less oxidative phosphorylation and glycolysis as a source of energy production, whereas wild-type efficiently used both oxidative phosphorylation and glycolysis for their ATP production (**Figure 6C**). In addition, KEGG-based oxidative phosphorylation analysis showed significant downregulation of ATP synthase gene activity in ST2-deficient macrophages (**Supplementary Figure 2**). These data suggest reduced macrophage energy expenditure in the absence of ST2 expression, especially during efferocytosis.

**Figure 6:**
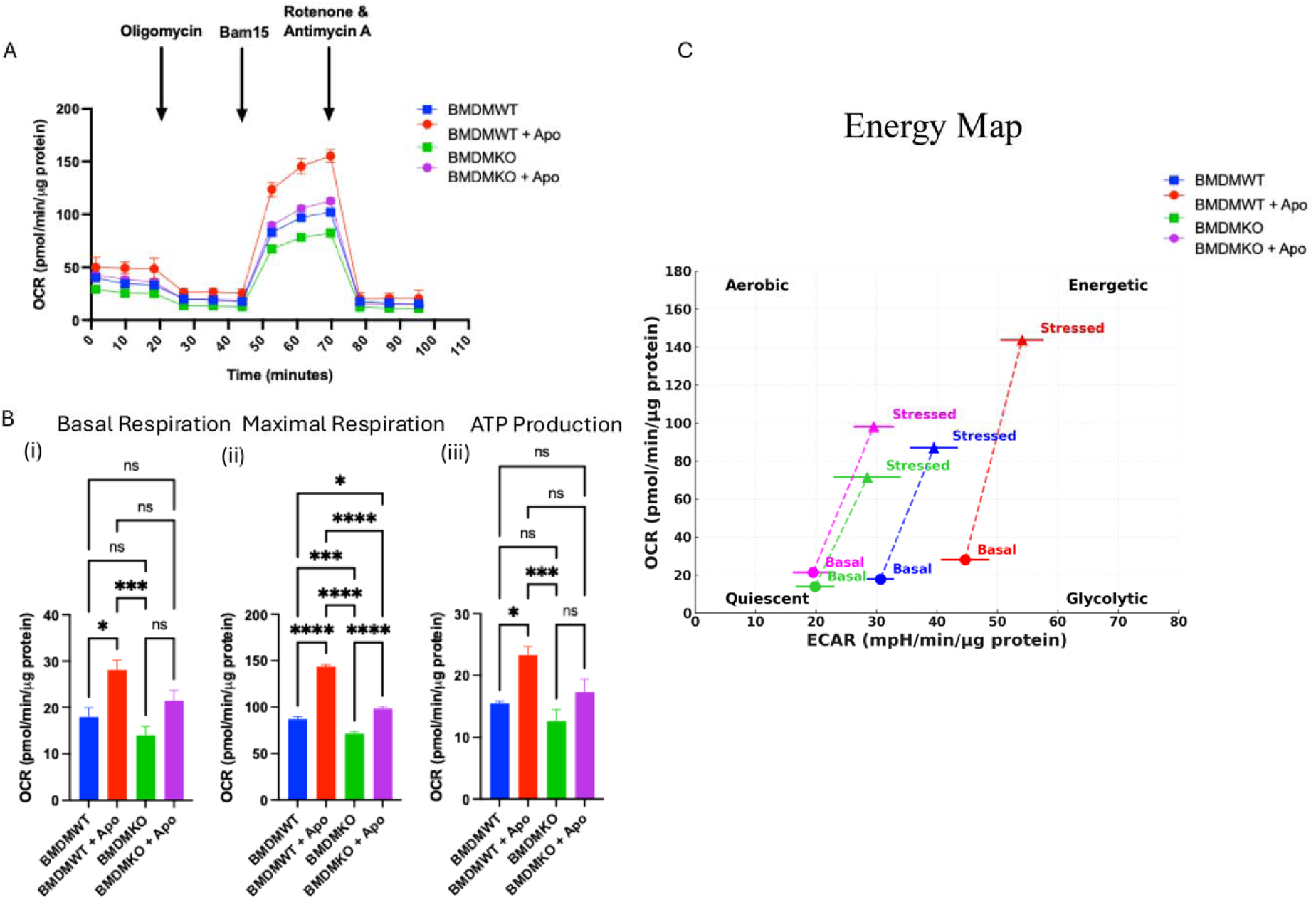
ST2 deficiency impairs the mitochondrial metabolism of macrophages. (A) Seahorse-based Cell mito stress assay of bone marrow-derived macrophages shows oxygen consumption rate (OCR) as a function of time under different metabolic stress conditions. (B) Quantitative analysis of (i) Basal Respiration, (ii) Maximal Respiration, and (iii) ATP Production highlights the metabolic differences between wild-type (WT) and ST2-deficient (ST2KO) macrophages under basal conditions and upon exposure to apoptotic cells (Apo). (C) Energy Map: OCR and extracellular acidification rate (ECAR) values from basal and stressed conditions are plotted, illustrating the distinct metabolic profiles of WT and ST2KO macrophages. WT macrophages (blue and red) show a greater shift towards high OCR and ECAR upon activation compared to ST2KO macrophages (green and magenta). The position of the basal and stressed conditions highlights the reduced metabolic flexibility and energetic capacity of ST2KO macrophages. Two replicates, n=3; mean±SEM is shown; *p<0.05; **p<0.01; ***p<0.001; ****p<0.0001.

## 4. Discussion

Innate immune cells such as macrophages are central not only in the pathogenesis of kidney disease but also exhibit therapeutic potential in limiting tissue injury and controlling fibrosis^49^. Macrophages and neutrophils are activated in response to AKI, producing cytokines and chemokines, causing inflammation, which if uncontrolled causes tissue damage and progression to CKD and end-stage renal disease (ESRD) among other causes^8,50^. Since AKI and CKD pose a significant burden to public health^51^, understanding the underlying molecular mechanisms is critical for the development of new therapeutic strategies. The study aimed to investigate the role of ST2 expression in myeloid cells on acute and chronic renal injury.

After ischemic injury, innate immune cells are activated and mobilized to the outer medulla where substantial tubular cell loss occurs^52^. Earlier studies have shown that damaged parenchymal cells release alarmins such as IL-33 to recruit innate immune cells^53^. IL-33 levels are elevated in the kidneys of the mice with AKI^54^. Multiple cell types including Tregs, ILC2, and macrophages can respond to the IL-33 released from damaged cells^20,30^. We have shown earlier that a bifunctional cytokine IL233, bearing the activities of IL-2 and IL-33, could increase Tregs and ILC2 to prevent and reverse ongoing AKI in multiple mouse models^28^. To understand the effect of IL-33 on Tregs, we recently investigated the role of ST2 expression on Tregs and their contribution to the repair of tubular injury in an AREG-dependent manner^30,55^. IL233 treatment also enhanced M2 macrophages and reversed ongoing chronic inflammation in mouse models of lupus nephritis and diabetic nephropathy^28,56^. The role of ST2 expression on macrophages remained a knowledge gap. Therefore, this study aimed to investigate the role of ST2 expression in myeloid cells on acute and chronic renal injury

We noted that myeloid cells, in particular macrophages, express ST2 in the kidneys of mice (**Figure 1**) and human macrophages also express ST2 (representative out of two human samples shown; **Supplementary Figure 3**). We addressed the role of ST2 in these cells through experiments with mice that lack ST2 expression in LysM-expressing myeloid cells (*ST2^fl/fl^ LysM^Cre^* mice). We demonstrated that ST2 deletion in myeloid cells can be beneficial during the acute bilateral IRI-AKI, but has deleterious consequences in the context of CKD, modeled by unilateral IRI followed by nephrectomy. These results correlate well with previous reports which showed that macrophage depletion at the acute phase of renal injury protects kidney function, whereas macrophage depletion at later time points after injury attenuates tubular proliferation and delays repair^18^. Previous studies have also suggested that macrophages present in the kidney during the acute phase of injury exhibit a pro-inflammatory phenotype akin to M1 macrophages^18^, which is consistent with our observations of the renal cytokine production of proinflammatory cytokines in acute injury settings (**Figure 1K**). On the other hand, during the later repair phase, macrophages appear to have an alternative activation-associated M2-like phenotype induced by IL-33^23^. Our study presented here suggests that the loss of the ST2 signaling axis in macrophages reduces the expression of M2-like signature markers on macrophages in chronic settings (**Supplementary Figure 6**) and a stronger signature of proinflammatory cytokines in the kidneys of mice with ST2-deficient macrophages (**Figure 2J**). These findings support a role for ST2 in the differentiation of macrophages to an M2-like state during chronic injury.

The M2 phenotype macrophages are known to induce immunological tolerance and support the differentiation of Tregs in acute tissue injury (reviewed in^57^). Interactions between innate and adaptive immune cells dictate the outcomes following injury^58^. Tregs play an important role in maintaining immune homeostasis and preventing excessive inflammation^59^. To study the interaction between macrophages and Tregs, we used a co-culture system of macrophages and naïve T cells and analyzed Treg differentiation and cytokine production profiles. These results showed that ST2 deficient macrophages co-cultured with responder T cells supported significantly reduced Treg differentiation and lower responder T-cell production of IL-4 and IL-10 with higher TNFα and IFNγ production. Even human macrophages when treated with IL-4 and IL-10, induce Tregs^60^. Such innate and adaptive immune interaction changes could also contribute to chronic inflammation in the context of CKD.

Through bulk RNA sequencing of BMDM, we demonstrated that the loss of ST2 expression significantly alters the transcriptional landscape and functional capacity of macrophages, highlighting its critical role in macrophage biology. Euclidean distance clustering and differential expression heatmaps revealed distinct transcriptional profiles in ST2-deficient macrophages. Unbiased differential gene expression analysis using a volcano plot identified several key genes associated with macrophage persistence, polarization, and tissue repair. These included *Bcl2a1b* (B-cell lymphoma 2-related protein A1b), critical for balancing cell survival and apoptosis^34^; *Arg1*, which promotes macrophage polarization toward the anti-inflammatory M2 phenotype^61^; *Gdf15*, a cytokine regulating inflammatory responses and tissue repair^62^; CD38, a multifunctional surface protein involved in immune activation and macrophage polarization^63^; and *Chil3* (Chitinase-like 3), a marker of M2 macrophages that contributes to anti-inflammatory environments and tissue repair^35^. A paralog of *Bcl2a1b*, *Bcl2l1* was also upregulated in macrophages in biopsies of patients with acute tubular necrosis^10^. It was shown by Lloyd Cantley group that ARG1 expression on macrophages is important for renal tubular proliferation and regeneration in renal IRI studies^64^. More recently, deletion of the immune activation-related phosphatase SHP2 in macrophages promoted M2 polarization and alleviated renal IRI^65^. Similar to the macrophage-specific deletion of ST2, *Gdf15* expression was also diminished in ST2-deficient Tregs as compared to ST2-sufficient Tregs, suggesting that IL-33/ST2 pathway may be involved in the induction of a reparative transcriptional program that includes expression of GDF15^30^ in multiple cell types including renal tubular cells and its deletion exacerbated tubular injury in renal IRI^66^. The upregulation of these genes in ST2-expressing macrophages suggests that ST2 signaling supports an anti-inflammatory and tissue-reparative macrophage phenotype. Pathway analysis further revealed that ST2 expression influences critical biological processes, including wound healing, metabolic pathways such as oxidative phosphorylation and glycolysis, chemokine responses, and phagosome formation. These findings indicate that ST2 signaling plays a central role in regulating macrophage-mediated immune modulation and tissue repair, underscoring its potential as a therapeutic target.

Macrophages are professional phagocytes with key roles in performing efferocytosis and clearance of dead cells, cellular debris, and pathogens^67^. We found increased numbers of uncleared dead cells in the kidneys of mice with ST2 deficiency in the myeloid lineage in both the acute and chronic models of renal injury, suggestive of defective efferocytosis *in vivo*. Uncleared cells and cell debris can lead to the persistence of proinflammatory signals, which can further exacerbate chronic inflammation. Importantly, *ex vivo* efferocytosis was significantly reduced in ST2-deficient macrophages, compared to control cells. Uncleared cells and cell debris can lead to the persistence of proinflammatory signals, which can further exacerbate chronic inflammation. The kidney resident macrophages were shown to upregulate receptors associated with the clearance of cell debris and immune complexes after renal IRI^11^. In addition, directed gene expression analysis of macrophage-specific functional markers demonstrated reduced expression of genes that encode proteins involved in promoting phagocytosis of dead/apoptotic cells (also known as ‘efferocytosis’), including *Mfge8*, *Itgb3*, *Abca1*, and *Stab2* in macrophages lacking ST2 (**Figure 5A**).

Since efferocytosis is an energy-consuming process, we also tested the role of ST2 in macrophage metabolism during efferocytosis using the Seahorse assays. M1 macrophages have an inhibition of the TCA cycle, whereas the M2 macrophages depend on constant ATP production via oxidative phosphorylation and fatty acid oxidation for the resolution of inflammation^68^. In agreement with a previous study^69^, we found that efferocytosis causes a metabolic shift in macrophages from initial glycolysis towards oxidative phosphorylation and increases ATP production. However, loss of ST2 was associated with reduced macrophage capacity to utilize oxidative phosphorylation, whereas control macrophages continued to utilize both oxidative phosphorylation and glycolysis for their ATP production, as noted by others^47,70^. These results suggest that ST2 plays an important role in regulating macrophage metabolic pathways, which could have implications for their function in the immune response. Whereas a previous report indicated that ST2^+^ airway macrophages were transcriptionally primed for support of epithelial repair^71^, our data indicate that ST2 also regulates macrophage cell cycle and activation. Since efferocytosis can stimulate macrophage proliferation to hasten tissue repair^48^, it will be important to discern whether the decreased proliferation of ST2 deficient macrophages is linked to reduced efferocytosis efficiency in our future studies.

In conclusion, our study identifies a crucial role of ST2 signaling in myeloid cells in the regulation of acute and chronic renal injury and highlights the impact of ST2 expression on macrophage homeostasis and function, including metabolism, proliferation, and efferocytosis. These findings provide valuable insights into the role of ST2 in regulating macrophages and its underlying effects on kidney injury.

## Data Sharing

The RNA-seq dataset reported in this study has been deposited in the Gene Expression Omnibus (GEO) under the accession number GSE288154 (Reviewer Access Token: mhghwaimdhulhqx). All data that support the findings are included within the manuscript. Additional supporting data are accessible in the supplementary file.

## Acknowledgments

We would like to acknowledge the University of Virginia (UVA) Research Histology Core, UVA Flow Cytometry Core, and UVA Genome Analysis and Technology Core for processing histology samples, FACS-sorting, and RNA-seq analysis, respectively.

## Author contributions

V.S. and R.S. designed the research; V.S., S.Ac., G.C., O.P. and B.M. performed research; C.U., N.L. and T.B., helped with Seahorse Metabolic analyses assays. S.Ar. performed Efferocytosis assays. V.S., S.Ar., and R.S. analyzed data; and V.S. and R.S. wrote the manuscript.

