## Supplementary Figures for "ST2 Signaling Regulates Innate Immune Responses in Kidney Injury"

**Supplementary Data and Information**

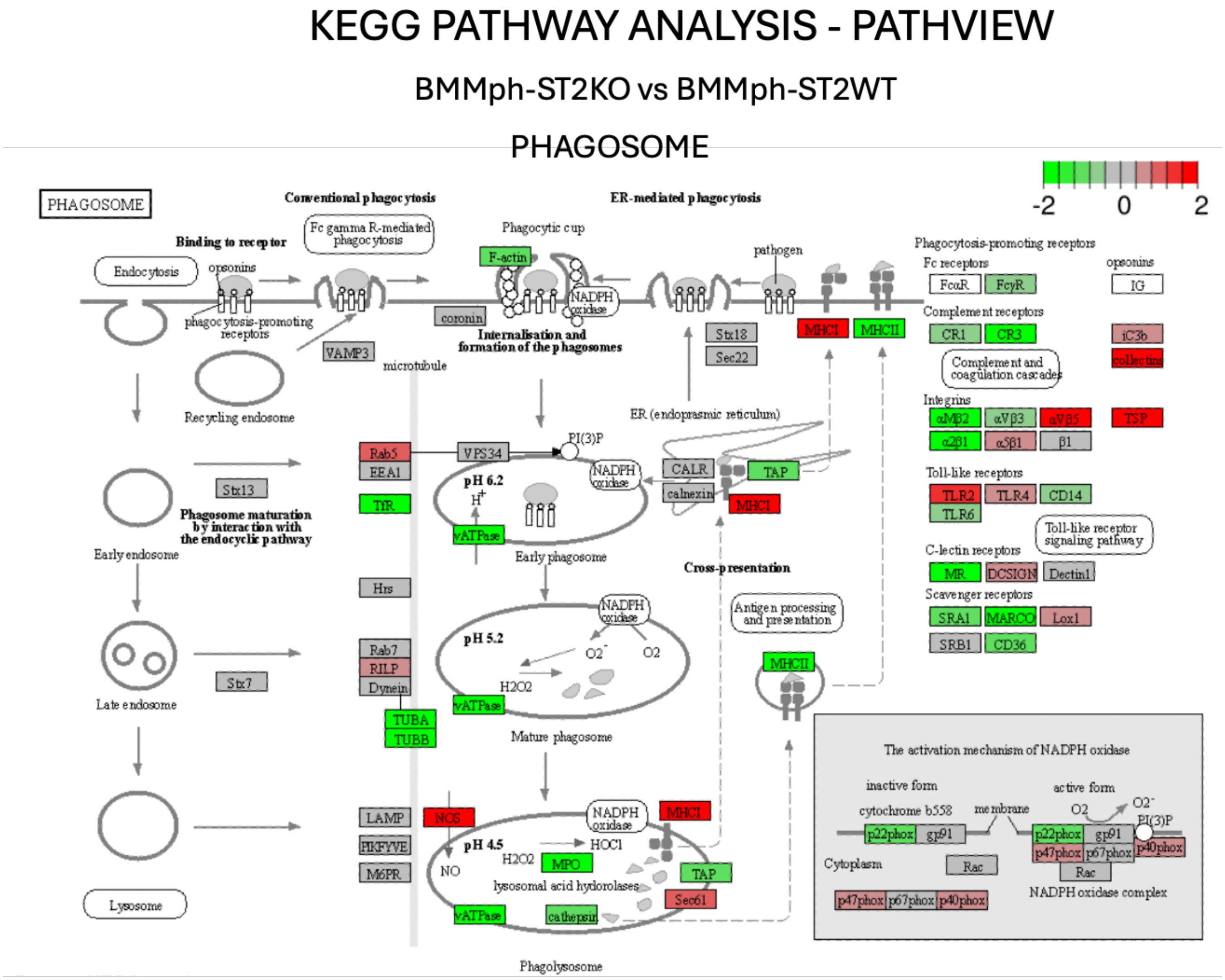


**Supplementary Figure 1:** KEGG-based phagosome pathway analysis indicating downregulation of critical genes BMMph ST2 knockout macrophages.

### KEGG PATHWAY ANALYSIS - PATHVIEW

#### BMMph-ST2KO vs BMMph-ST2WT

#### OXIDATIVE PHOSPHORYLATION

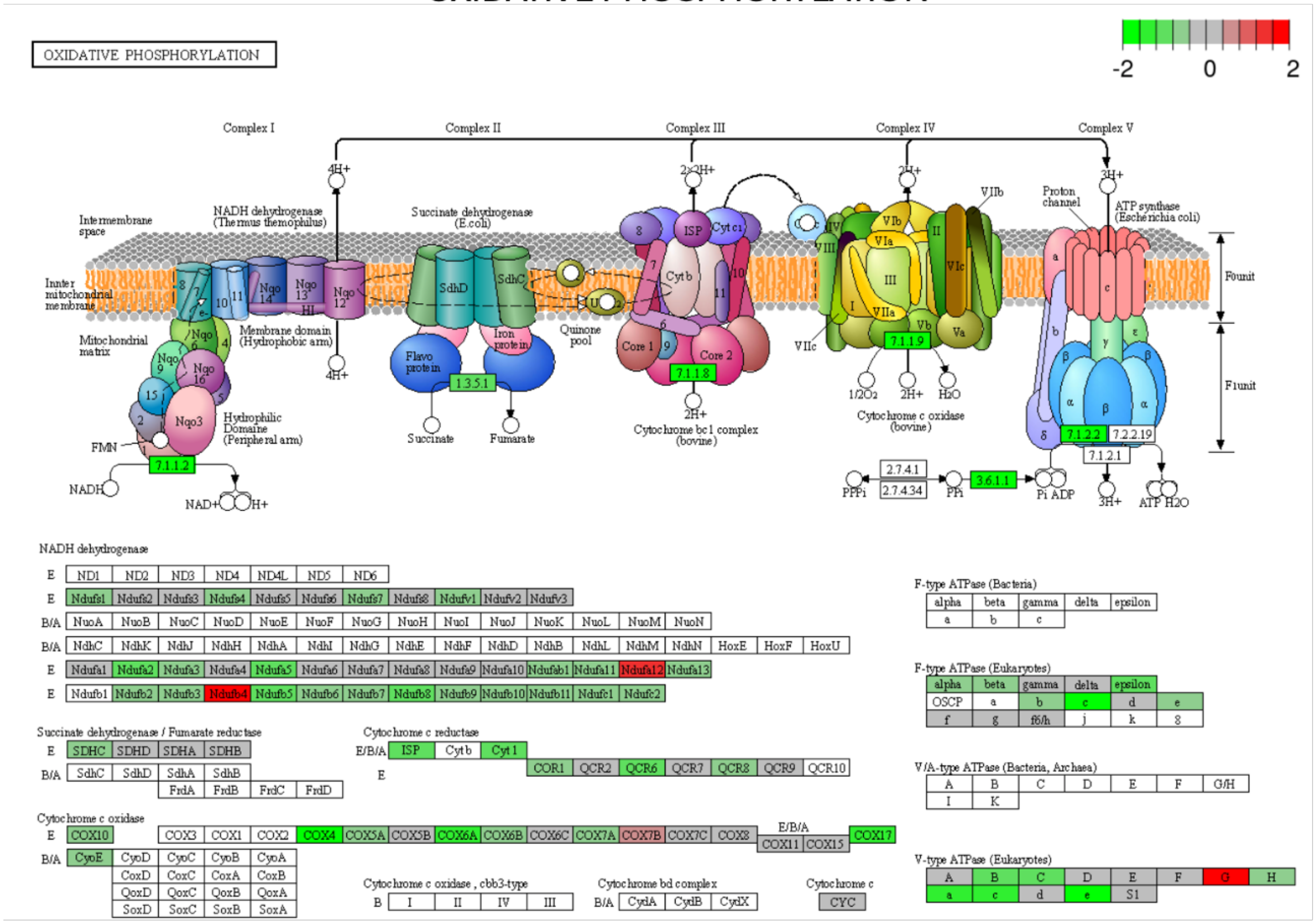


**Supplementary Figure 2:** KEGG-based phagosome pathway analysis indicates the downregulation of critical genes involved in ATP production in BMMph ST2 knockout macrophages.

Expression of ST2 on human Tonsillar CD11b+CD11c+ cells

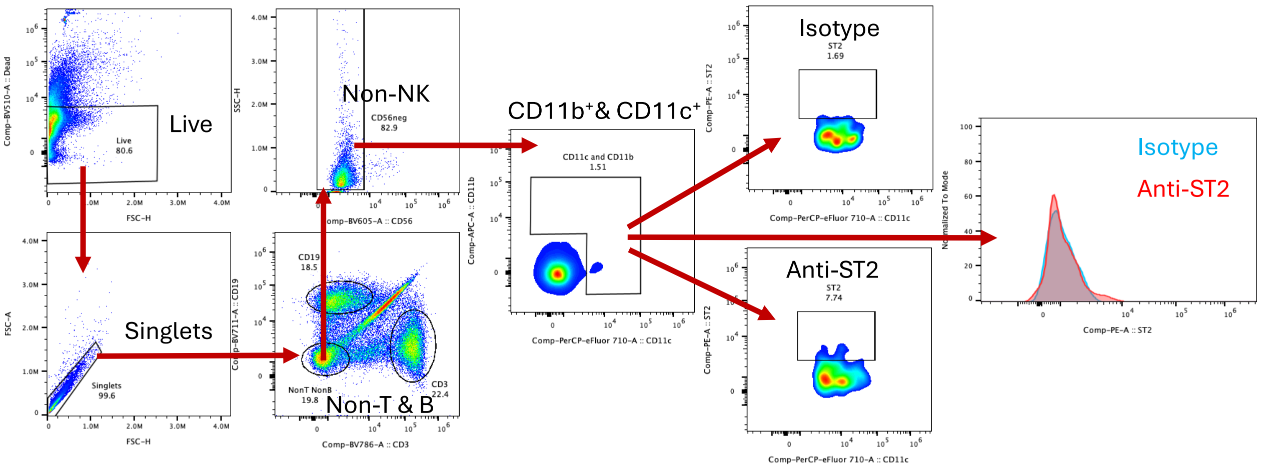


**Supplementary Figure 3:** Expression of ST2 on Human Tonsillar CD11b$^{+}$CD11c$^{+}$ cells. One representative out of two human tonsil samples shown.

Kidney Injury and Fibrosis Markers – Acute Injury

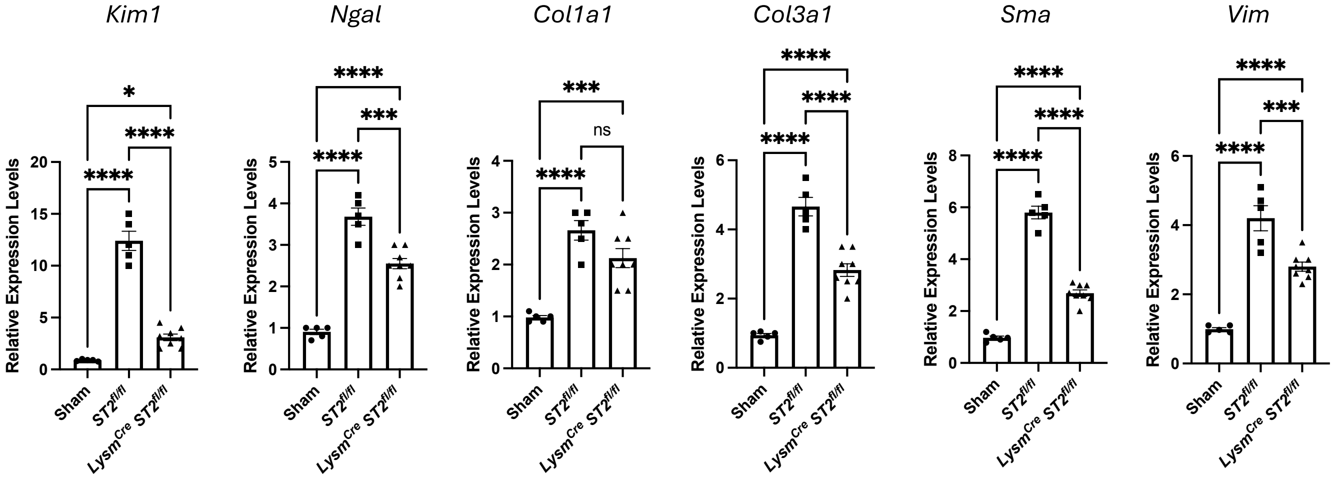


**Supplementary Figure 4:** Quantitative expression analysis of kidney Injury and fibrosis markers following acute kidney injury (AKI). Symbols represent individual mice (n>5); mean±SEM is shown; *p<0.05; ***p<0.001; ****p<0.0001.

Kidney Injury and Fibrosis Markers – Chronic Injury

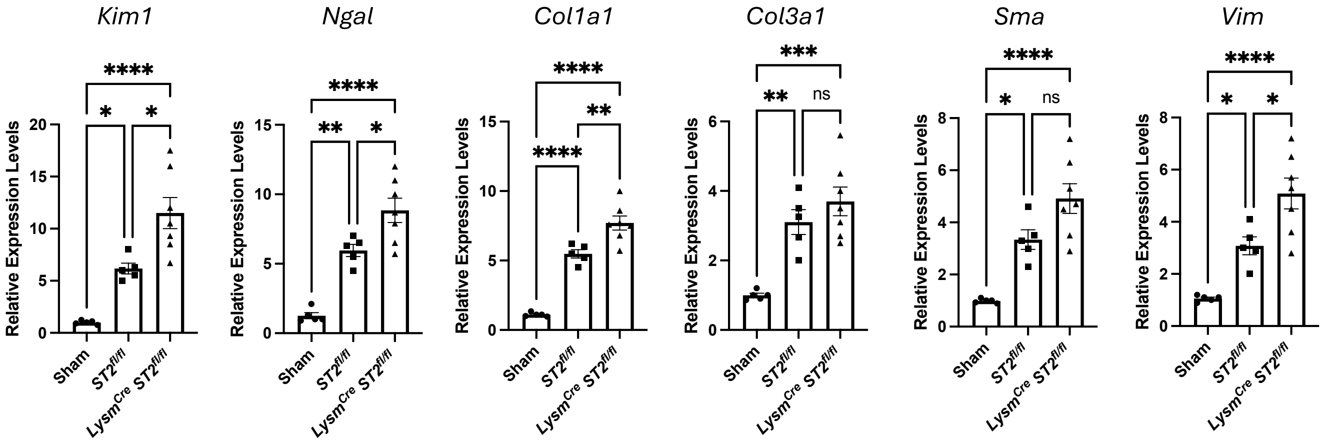


**Supplementary Figure 5:** Quantitative expression analysis of kidney Injury and fibrosis markers following chronic injury (CKD). Symbols represent individual mice (n>5); mean±SEM is shown; *p<0.05; **p<0.01; ***p<0.001; ****p<0.0001.

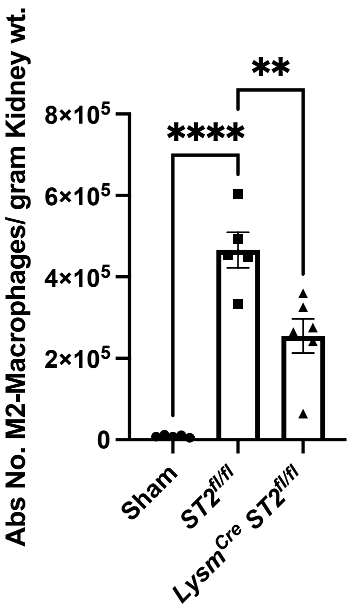


**Supplementary Figure 6:** Flow cytometry-based quantification of CD206$^{+}$ M2 Macrophages in Chronic Injury (CKD). Symbols represent individual mice (n>5); mean±SEM is shown; **p<0.01; ****p<0.0001.

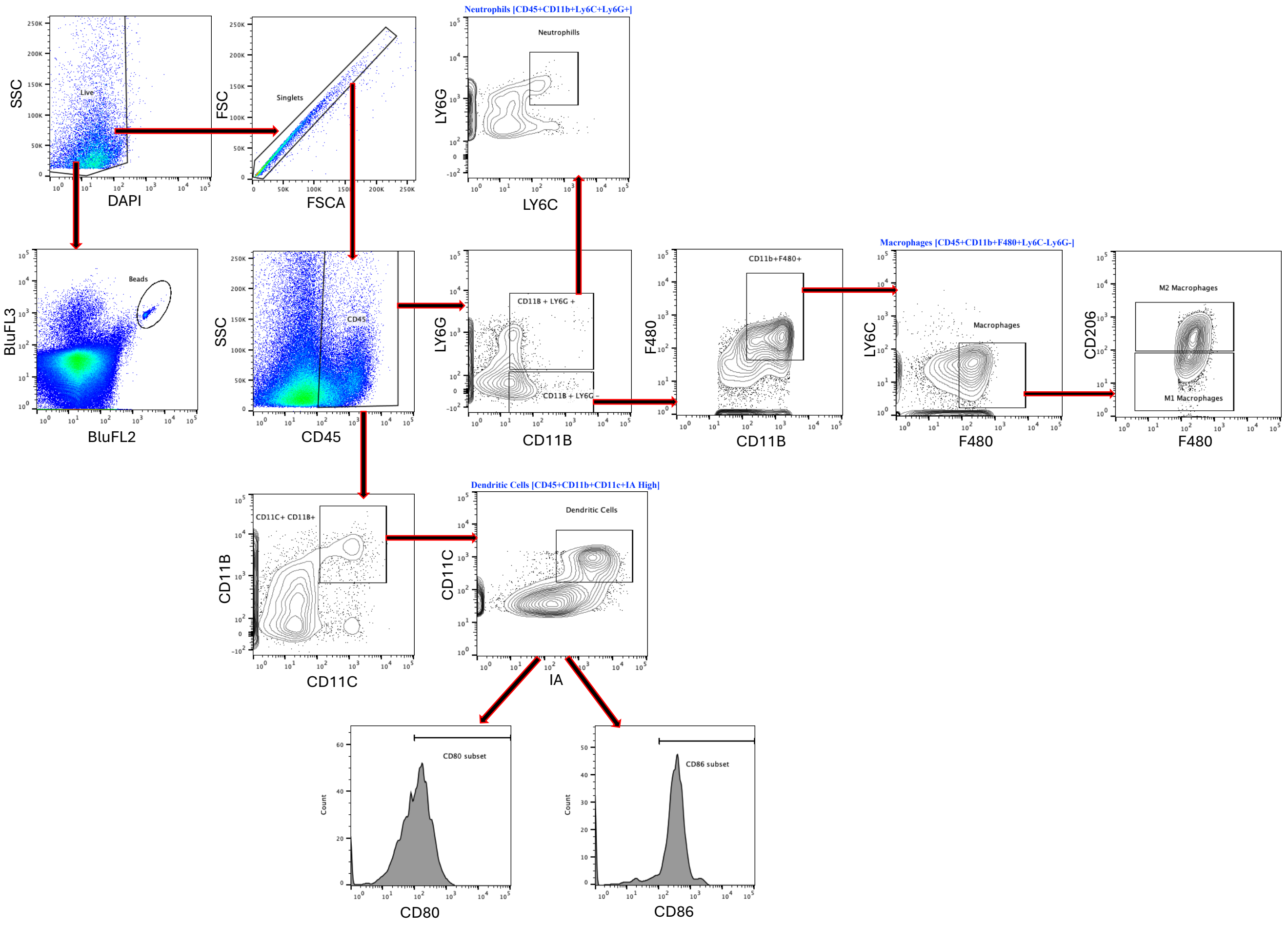


**Supplementary Figure 7:** Flow cytometry gating strategy for Neutrophils, Macrophages, and Dendritic cells in kidneys.

**Supplementary Table 1: Antibodies Used for Flow Cytometry/Immunostaining**

| Antigen | Species | Fluorophore | Catalog No. | Vendor |
|---|---|---|---|---|
| Ly6G | Mouse | BV605 | 108440 | BioLegend |
| Ly6C | Mouse | PerCP-Cy5.5 | 128012 | BioLegend |
| CD45 | Mouse | PerCP | 103130 | BioLegend |
| CD11b | Mouse | APC-e780 | 47-0112-82 | Life Technologies |
| CD11c | Mouse | BV570 | 117331 | BioLegend |
| F480 | Mouse | Alexa647 | 123122 | BioLegend |
| CD206 | Mouse | BV711 | 141727 | BioLegend |
| IA | Mouse | PerCP-Alexa710 | 46-5321-80 | Life Technologies |
| CD80 | Mouse | FITC | 553768 | BD |
| CD86 | Mouse | PE | 12-0862-82 | Life Technologies |
| INFγ | Mouse | PerCP-Cy5.5 | 505822 | BioLegend |
| IL-4 | Mouse | BV421 | 504127 | BioLegend |
| TNFα | Mouse | APC-Cy7 | 506344 | BioLegend |
| IL-10 | Mouse | APC | 505010 | BioLegend |
| CD3 | Mouse | BV750 | 100249 | BioLegend |
| CD4 | Mouse | BV650 | 100469 | BioLegend |
| CD8 | Mouse | PE-Cy5 | 100710 | BioLegend |
| ST2 | Mouse | PE | 566311 | BD |
| Foxp3 | Mouse | BV421 | 562996 | BD |
| IL-17 | Mouse | PE-Cy7 | 506922 | BioLegend |
| CD56 | Human | BV605 | 740405 | BD |
| CD3 | Human | BV785 | 317330 | BioLegend |
| CD11c | Human | PerCP-eFlour-710 | 46-0116-41 | Life Technologies |
| CD11b | Mouse/Human | APC | 101212 | BioLegend |
| ST2 | Human | PE | 12-9338-42 | Life Technologies |

**Supplementary Table 2: Real-time PCR Primers**

| **Primers** | **Sequence 5'-3'** |
|---|---|
| *Kim1* Fwd | ACATATCGTGGAATCACAACGAC |
| *Kim1* Rev | ACTGCTCTTCTGATAGGTGACA |
| *Ngal* Fwd | TGGCCCTGAGTGTCATGTG |
| *Ngal* Rev | CTCTTGTAGCTCATAGATGGTGC |
| *Col1a1* Fwd | GCTCTTTTTAGATACTGTGGTGAGGAA |
| *Col1a1* Rev | GTTTCCACGTCTCACCATTG |
| *Col3a1* Fwd | ACAGCTGGTGAACCTGGAAG |
| *Col3a1* Rev | ACCAGGAGATCCATCTCGAC |
| *Vimentin* Fwd | GATCGATGTGGACGTTTCCAA |
| *Vimentin* Rev | ATACTGCTGGCGCACATCAC |
| *Acta2* Fwd | CTGACAGAGGCACCACTGAA |
| *Acta2* Rev | AGAGGCATAGAGGGACAGCA |

**Supplementary Table 3: Genotyping Primers**

| **Primers** | **Sequence 5'-3'** |
|---|---|
| IL1RL1 Flox Fwd | CCATACTGTGAGATGGCGC |
| IL1RL1 Flox/WT Rev | GAAGGAGGAAATTCAGACTGGG |
| IL1RL1 Fwd | GGCCACCAGATCATCACAGTTGAAGG |
| ST2 WT Fwd | GCTGGATAAAGCTATATCATGG |
| ST2 KO Fwd | GATTGCACGCAGGTTCTC |
| ST2 WT/KO Rev | GGAAATGCAACCAGAAGTGCACAGG |
| Generic Cre Fwd | AGGTGTAGAGAAGGCACTTAGTAGC |
| Generic Cre Rev | CTAATCGCCATCTTCCAGCAGG |
